# Detection, not mortality, constrains the evolution of virulence

**DOI:** 10.1101/2021.11.14.468516

**Authors:** David A. Kennedy

## Abstract

Why would a pathogen evolve to kill its hosts when killing a host ends a pathogen’s own opportunity for transmission? A vast body of scientific literature has attempted to answer this question using “trade-off theory,” which posits that host mortality persists due to its cost being balanced by benefits of other traits that correlate with host mortality. The most commonly invoked trade-off is the mortality-transmission trade-off, where increasingly harmful pathogens are assumed to transmit at higher rates from hosts while the hosts are alive, but the pathogens truncate their infectious period by killing their hosts. Here I show that costs of mortality are too small to plausibly constrain the evolution of disease severity except in systems where survival is rare. I alternatively propose that disease severity can be much more readily constrained by a cost of behavioral change due to the detection of infection, whereby increasingly harmful pathogens have increasing likelihood of detection and behavioral change following detection, thereby limiting opportunities for transmission. Using a mathematical model, I show the conditions under which detection can limit disease severity. Ultimately, this argument may explain why empirical support for trade-off theory has been limited and mixed.

## Introduction

The seminal works of Anderson and May (Anderson and May 1982; May and Anderson 1983) changed the way that biologists thought about the evolution of pathogen virulence, defined as the severity of disease signs or symptoms caused by infection with a particular pathogen. Before Anderson and May, the conventional wisdom was that pathogens would evolve to be avirulent over time (Alizon and Michalakis 2015), since a highly virulent pathogen risks killing its host and by killing its host a pathogen truncates its own infectious period and reduces its own fitness. Anderson and May articulated that natural selection favors pathogen variants that maximize their own fitness. If virulence were correlated with other epidemiological parameters such as infectiousness or time to recovery, intermediate levels of virulence could maximize fitness, and thus be evolutionarily adaptive. The idea they proposed, “trade-off theory”, is that the cost of virulence, which they assumed was a truncated duration of infectiousness caused by host mortality, trades off against other benefits such as an increased rate of transmission or a decreased rate of recovery. This work has been hugely influential, and the trade-off theory that they proposed has since been termed the “new conventional wisdom” (Alizon et al. 2009). Ultimately, trade-off theory was meant to explain why evolution has generated pathogens that have intermediate levels of virulence. That is, 1) why do pathogens harm their hosts at all, and 2) why do they not harm their hosts more?

There are only a limited number of explanations for why evolution has allowed pathogens to maintain virulence. Either there is no genetic variation for reduced virulence, selection is too weak to eliminate virulence entirely, or virulence is associated with some fitness benefit to the pathogen (Alizon and Michalakis 2015). Although it is possible to point to specific examples where each of these explanations might apply, the widespread detection of variation in virulence (e.g., Froissart et al. 2010) and the observation that virulence often increases during serial passage experiments (Ebert 1998) challenge the generality of these first two explanations. In contrast, a recent meta-analysis of experimental studies on virulence evolution found that replication rates within hosts positively correlates with both transmission potential and virulence (Acevedo et al. 2019), lending support to the third explanation. It has thus been generally accepted that, for most pathogens, some degree of virulence provides or is correlated with a fitness benefit in some environment where selection is acting.

This leaves the question of why pathogens are not more harmful to their hosts. Classical trade-off theory proposes that pathogens are not more harmful to their hosts because the fitness benefits associated with increased virulence saturate relative to the fitness costs of increased virulence (Alizon et al. 2009). Translated to a mathematical framework, the typical assumption is that increases in transmission rate saturate relative to increases in host mortality (Fig. 1). Such a relationship may emerge due to within host processes (Alizon and van Baalen 2005), and has been seen in some biological systems (e.g. De Roode et al. 2008; Read et al. 2015), but this saturation has not been generally detected (Acevedo et al. 2019). In fact, experimental data in support of trade-off theory has been disappointingly limited (Bull 1994; Lipsitch and Moxon 1997; Alizon et al. 2009; Alizon and Michalakis 2015; Cressler et al. 2016), leading to questions about the usefulness of trade-off theory entirely (Lipsitch and Moxon 1997; Ebert and Bull 2003; Bull and Lauring 2014).

**Figure 1:**
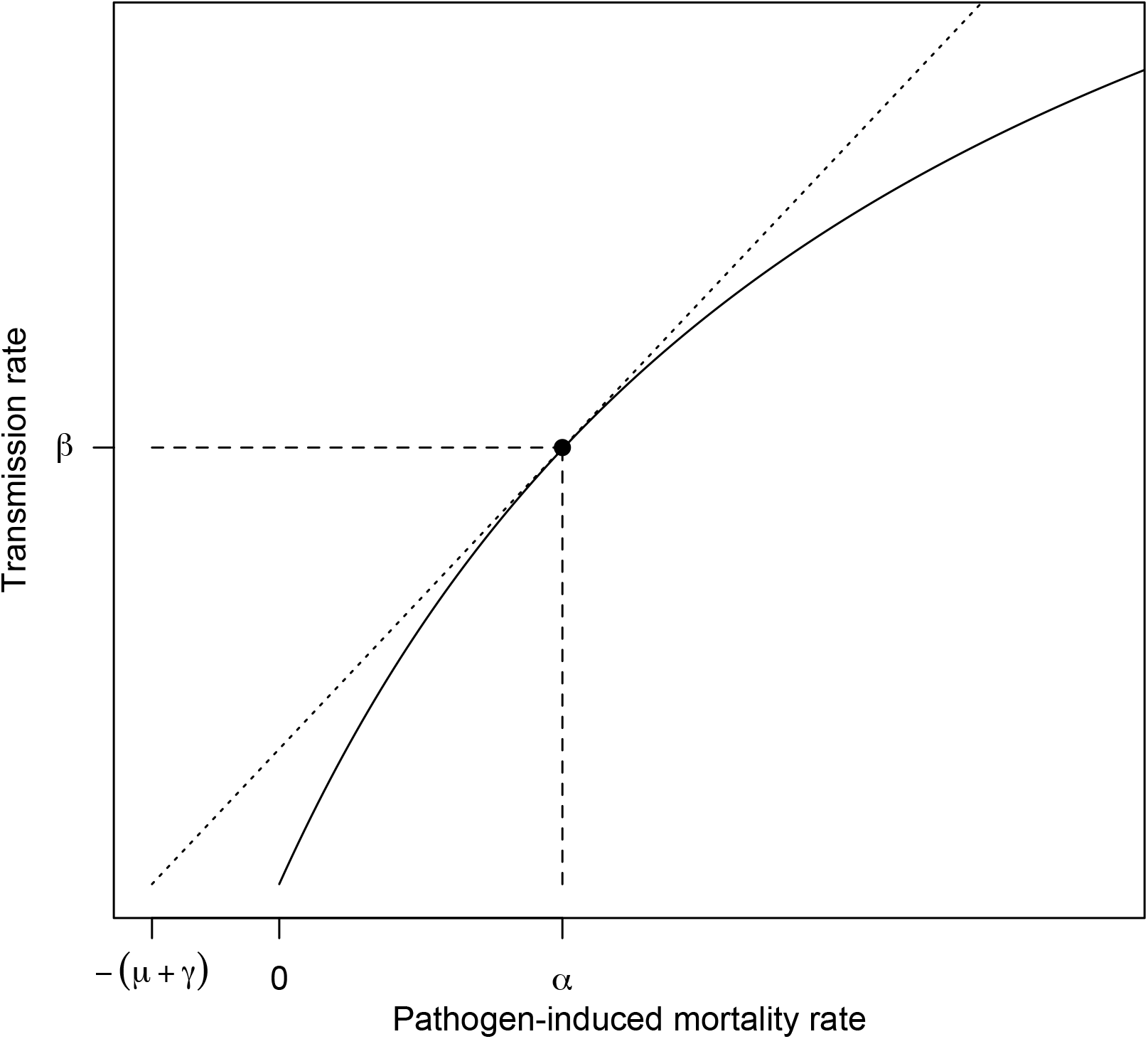
Classical formulation of the virulence-transmission trade-off. The solid curve shows a possible trade-off between transmission rate and pathogen induced host mortality rate. The evolutionarily optimal values of transmission rate *β* and mortality rate *α* according to original theory are depicted by the point where the dotted line touches the curve (Alizon et al. 2009).

Rather than disregard trade-off theory, some have argued that the lack of experimental support for trade-off theory has resulted from difficulties in designing appropriate experiments (Alizon and Michalakis 2015) or from collecting inappropriate proxies for virulence and transmission (Cressler et al. 2016). I argue that the reason few experiments have found evidence supporting trade-off theory is that they assume the cost of virulence is borne out through a reduction in the duration of infections due to host mortality, despite the fact that this is rarely the case.

Behavior and behavioral changes are being increasingly recognized as important drivers of infectious disease dynamics in humans (Funk et al. 2010) and other animals (Stockmaier et al. 2021). Changes in behavior alone are capable of tipping the balance from localized pathogen extinction to successful disease emergence (Alexander and McNutt 2010; Shaw and Kennedy 2021). Pathogen-induced changes in behavior could therefore impose substantial selection pressure. For example, if contact rates between hosts declined with increasing disease severity, there could be strong selection pressure on the pathogen to reduce its severity. Such a relationship between disease severity and contact rates has been observed (McKay et al. 2020), yet with few exceptions (e.g. Ewald 1983, 1994), the role of behavior on virulence evolution has been largely ignored. The fact that disease ecology is rarely driven by host mortality but can often be driven by host behavior might lead one to wonder whether changes in behavior following infection are generally a stronger evolutionary force than changes in mortality.

Here I argue that the cost of virulence typically plays out through morbidity-induced reductions in contact rates – which I refer to as a “detection cost of virulence” – and that this cost of virulence drastically outweighs the cost of infection induced mortality for the vast majority of systems. Note that this argument builds on several previously published concepts. Ewald (1983, 1994) long ago proposed a trade-off between virulence and transmission mode that implicitly included a virulence-detection trade-off. This idea was later formalized (Day 2001, 2002a), and the concept has been discussed by many others (for example, Alizon and Michalakis 2015; McKay et al. 2020). Likewise, Ebert and Bull (2003) and Bull and Lauring (2014) previously discussed that virulence, in the context of mortality, is likely to impose only an indirect and weak evolutionary cost.

Here I show that, under the assumptions of a mortality-rate-transmission-rate trade-off, the cost of virulence can be written in terms of the infection fatality rate (defined as the fraction of all infections, symptomatic and asymptomatic, that result in disease-induced host death). Using this new form, I show that mortality is too weak a cost to constrain the virulence of pathogens in most systems, but that detection costs can.

## Model and Results

### The cost of mortality

The original formulation of the virulence-transmission trade-off arises from analysis of a classic SIR model based on the models of Kermack and McKendrick (1991) and Anderson and May (1979).

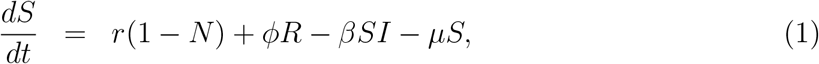

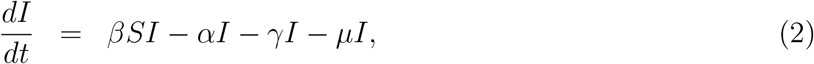

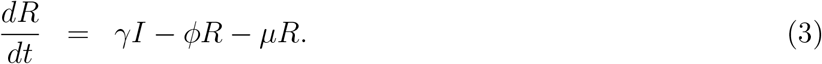

Above, *S, I*, and *R* are the respective densities of susceptible, infectious, and recovered hosts. *N* is the total population density derived by summing *S, I*, and *R. r* is the maximum per capita birth rate, *ϕ* is that rate at which immunity wanes, *β* is the transmission rate, *γ* is the recovery rate, *µ* is the baseline host mortality rate, and *α* is the pathogen-induced host mortality rate.

Under the assumptions of this model and any model that excludes non-linear environmental feedbacks such as spatial structure (Boots et al. 2004; Berngruber et al. 2015), coinfection (May and Nowak 1995), superinfection (Nowak and May 1994), host heterogeneity (Regoes et al. 2000), and non-linear transmission (Lion and Metz 2018), a pathogen strain that maximizes the basic reproductive number *R*_0_ will competitively exclude all other pathogen strains once the system reaches an equilibrium. It thus follows that natural selection will lead to the evolution of a pathogen strain that maximizes *R*_0_ (Anderson and May 1982).

In the above model, the basic reproductive number is

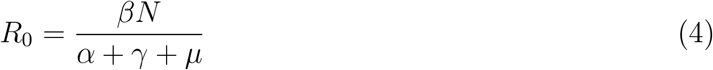

This formulation of *R*_0_ illustrates the paradox of virulence pointed out by May and Anderson (1983). That is, all else equal, a strain with lower virulence (i.e. smaller *α*) would have a higher *R*_0_, and thus, pathogens should evolve to be avirulent. However, if transmission rate *β* or recovery rate *γ* were functions of virulence, it need not be the case that low virulence is always favored. Famously, *R*_0_ can be maximized at intermediate virulence if the transmission rate *β* is a saturating function of host induced mortality *α* (Fig. 1). This so called virulence-transmission trade-off is by far the most widely invoked explanation for the maintenance of virulence in nature.

According to the principle of *R*_0_ maximization, a new pathogen variant would be able to invade and displace an existing pathogen strain provided the new value of *R*_0_ is greater than the old value of *R*_0_. Under the assumption that recovery rate *γ* is the same for two pathogen variants, this can be reduced to (Supplemental Information):

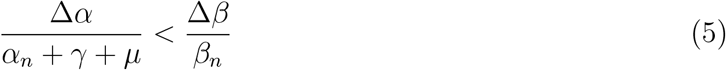

Above, I use the symbol Δ as shorthand for the difference between the old and new values for a parameter, such that Δ*X* corresponds to *X*_*n*_ *− X*_*o*_, where subscript “*n*” denotes the new variant and subscript “*o*” denotes the old variant. Inequality 5 leads to the well known result that if the transmission rate *β* is a saturating function of disease induced mortality rate *α*, then an optimal level of virulence can be derived as shown in Fig 1.

In a classical SIR model such as that described by Eq. 1-3 the infection fatality rate *F*, defined as the fraction of all infections that result in disease-induced death, can be written as 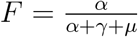. Note that the infection fatality rate is similar to the case fatality rate except that the latter often excludes asymptomatic infections. Under the assumption that recovery rates do not differ between variants (Day 2002b), Inequality 5 can be rewritten in terms of *F*, leading to the conclusion that a new mutation will spread if (Supplemental Information):

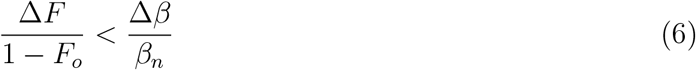

The above inequality states that a new variant will be able to invade and displace the current pathogen if the percentage decrease in infection survival rate 1 *− F*, is less than the percentage increase in the transmission rate *β*. Note that the left side of this inequality can be viewed as the costs of virulence and the right side can be viewed as the benefits.

The advantage of Inequality 6 over the standard formulation (Inequality 5) is that it shows that the cost of mortality depends on percent changes in survival rather than percent changes in mortality. For a pathogen with low survival (i.e. *F*_*o*_ *≈* 1), small changes in mortality can thus generate large constraints, but for pathogens with high survival (i.e. *F*_*o*_ *≈* 0), small changes in mortality generate small constraints since the denominator on the left hand side is approximately 1. The consequence of this asymmetry means that for pathogens with initially low infection fatality rates, almost all variants with increased transmission and virulence should be able to invade and spread. The fact that low virulence pathogens retain low virulence, however, suggests that there is something wrong with the theory.

To illustrate this point, consider a theoretical pathogen with a low infection fatality rate *F*_*o*_ *≈* 0, something akin to a rhinovirus that causes the common cold. If this pathogen has an *R*_0_ of 5 and an infection duration of 5 days, then that implies each infection produces 1 new infection per day. Inequality 6 tells us that a mutation that increased its per day infectiousness from 1.00 to 1.01 would be evolutionarily favored provided it did not increased the infection fatality rate above approximately 1%. Notably, a 1% change in transmission is small relative to differences in transmission rates typically detected between field isolates (e.g. Mackinnon and Read 1999), but this change in mortality rate is larger than the difference between a common-cold-causing rhinovirus and SARS-CoV-2 (O’Driscoll et al. 2021). Theory thus predicts that if the main cost of virulence were host mortality, the common cold could become as severe as COVID-19 in exchange for a 1% increase in the transmission rate of the virus. Yet no such variant has ever spread, and there has never even been a documented cluster of rhinovirus infections with COVID-like mortality rates. Similarly, an increase in transmission rate from 1.00 to 1.10 could justify an infection fatality rate as high as 9%, which is comparable to the infection fatality rate of the 2003 SARS virus (Parry 2003). Nearly identical numbers can be derived for pathogens and parasites that are typically thought of as moderately virulent, such as influenza A viruses, measles virus, *Plasmodium falciparum*, and SARS-CoV-2 (Fig. 2).

**Figure 2:**
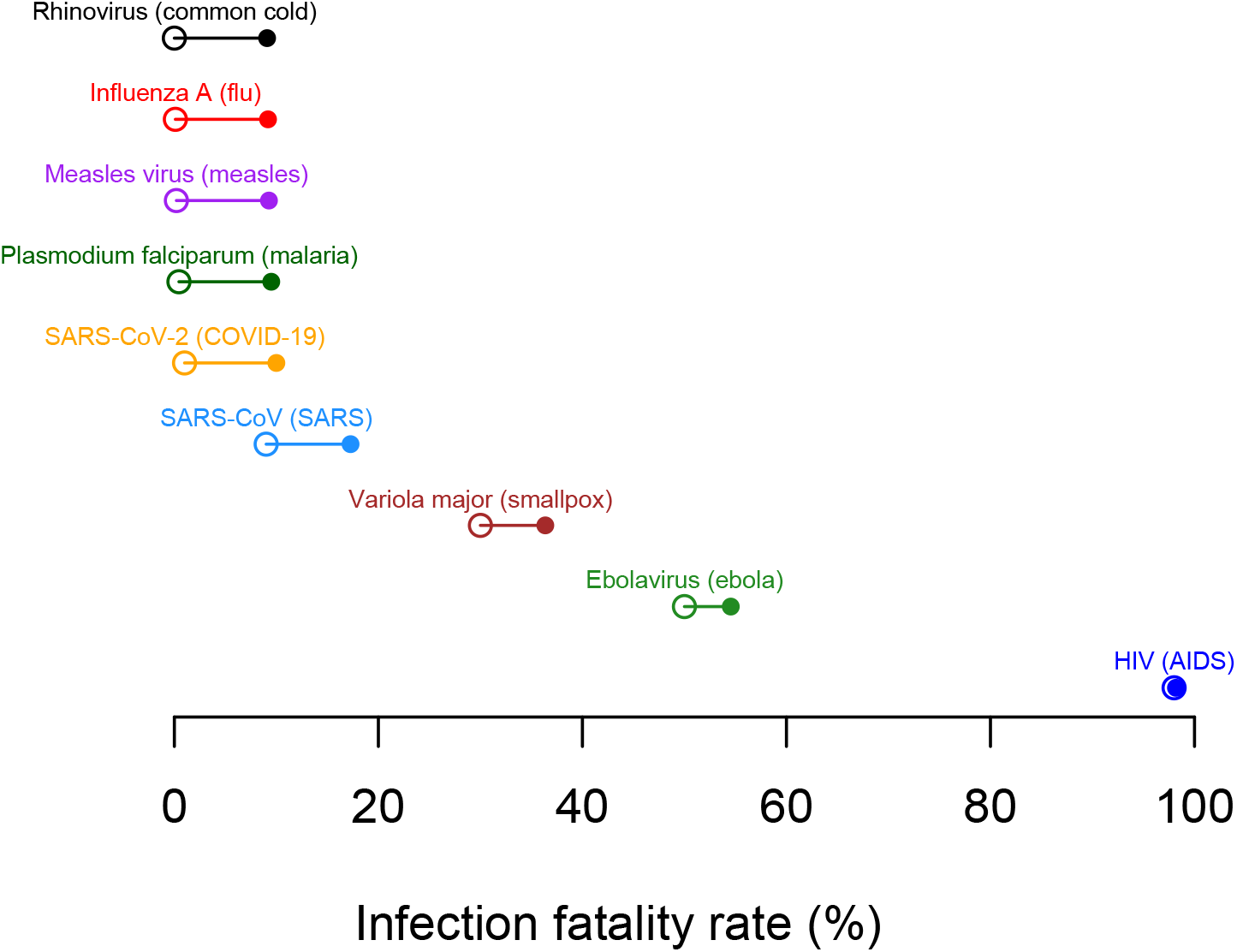
Under the assumption that host mortality constrains virulence, moderate changes in transmission rates can justify large increases in the infection fatality rate. Open circles indicate approximate infection fatality rates for various pathogens and parasites (values are for illustration purposes and may not be exact). Filled circles indicate the maximum infection fatality rate that would be evolutionarily favored under current theory if it were accompanied by a 10% increase in the transmission rate 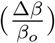 Note that this change in transmission is enough justify an otherwise harmless pathogen evolving to become as virulent as the 2003 SARS virus.

Under the above theory, mortality only provides a strong cost of virulence to pathogens with extremely high infection fatality rate *F* (Fig. 3) such as for lethal, chronic infections like human immunodeficiency virus (HIV, Fraser et al. 2014). One is therefore left to wonder what typically constrains disease severity if not infection-induced mortality.

**Figure 3:**
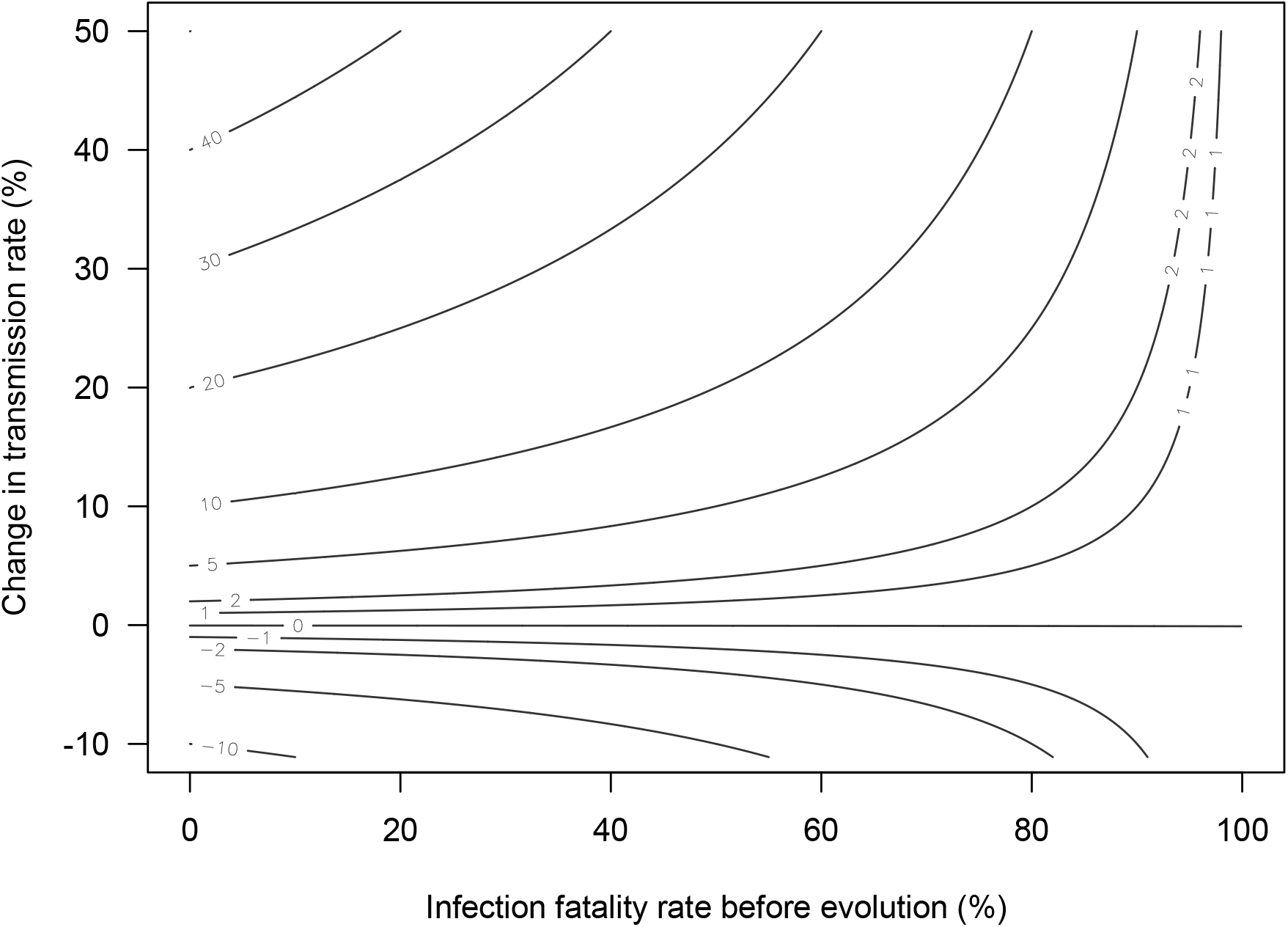
Contour lines show the maximum absolute change in the infection fatality rate that would be evolutionarily favored for a given percent change in transmission rate (i.e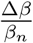). Note that when the original infection fatality rate is small (i.e. small values on x-axis) any absolute change in the infection fatality rate can be fully balanced by an equivalent percentage change in the transmission rate. The horizontal nature of the contour lines at small to moderate x-axis values indicates that costs of mortality are small unless the original infection fatality rate is large. Thus it is only when infection fatality rates before evolution are large, that increases in infection fatality pose a strong constraint on pathogen evolution.

## The cost of detection

I propose that a more reasonable mechanism constraining virulence is a reduction in transmission attributable to behavioral changes that follow detection of infection. For instance, once someone realizes they have COVID-19 symptoms, they may self-isolate thereby reducing transmission opportunities. Alternatively, an infected host may simply feel too sick to conduct their normal daily activities such as attend school or go to work, again reducing transmission opportunities. In either case, causing detectable infection would negatively impact pathogen fitness, and presumably, moreso for increasingly severe disease.

To formalize this concept, consider an alternative SIR-type model, with modification from Eqs 1-3. The differences are that 1) host mortality has been removed, and 2) the infectious class has been split up into three groups.

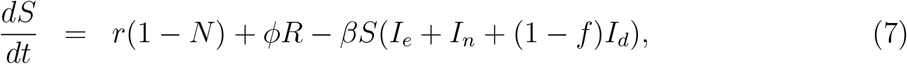

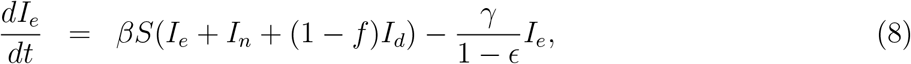

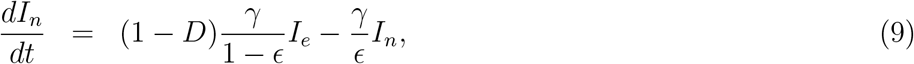

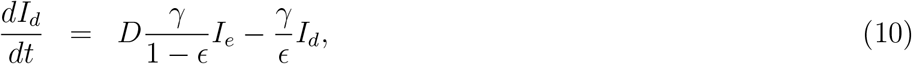

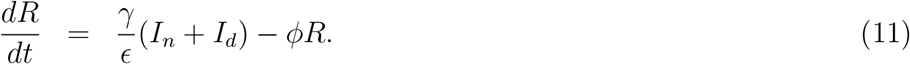

As before, *N* is the sum of host density in all classes, *S* is the density of susceptible hosts, and *R* is the density of recovered hosts. *I*_*e*_ tracks hosts in the early phase of infection in which detection of infection is not yet possible, and *I*_*n*_ and *I*_*d*_ track hosts in the later phase of infection for respectively not detected and detected infections. As before, the parameter *ϕ* is the rate at which immunity wanes, *γ* is the rate of recovery of infected individuals, and *β* is the transmission rate in the absence of detection. The parameter *f* is the reduction in transmission of detected infections relative to non-detected infections, *ϵ* is the fraction of an infection’s duration that occurs after detection is possible, and *D* is the fraction of infections that are detected. Note that the new parameters *f, ϵ*, and *D* are bounded between 0 and 1, while the reused parameters *ϕ, β* and *γ* can still take any non-negative value.

As before, we can readily derive the basic reproductive number,

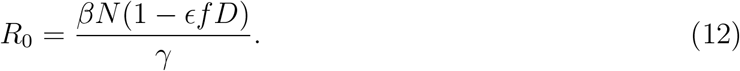

This formulation of *R*_0_ demonstrates the cost of detection as described by the parameter combination *ϵf D*. Note that this parameter combination describes the fraction of new infections that are avoided because of pathogen detection. It can be conceptualized as the fractional reduction in transmission that occurs over the lifetime of an infection due to detection. This may be realized through, for example, a reduction in contact rate or a reduction in infectiousness given contact. More highly virulent pathogens presumably lead to detection in a larger fraction of hosts (i.e. increased *D*), and more stringent actions to reduce transmission once detected (i.e. increased *f*) (McKay et al. 2020). This effect is only borne out for *ϵ*, the fraction of each infection that occurs after detection is possible. The net effect of these morbidity effects is to decrease overall transmission opportunities and thus *R*_0_. If we assume that increased disease severity correlates with increased transmission potential in the absence of detection *β*, as appears to be the case (Acevedo et al. 2019), then it is possible for *R*_0_ to be maximized at intermediate levels.

As before, a new pathogen variant would be able to displace an existing pathogen if the new value of *R*_0_ is greater than the old value of *R*_0_. As with a mortality cost, I assume the recovery rate *γ* is unchanged by evolution of virulence (i.e. no trade-off between virulence and recovery). If *ϵ*, the fraction of an infection that occurs after detection is possible, is also unchanged by evolution, then a new variant will be capable of invading if (Supplemental Information):

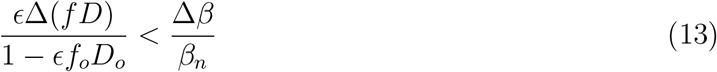

Above, *ϵ*Δ(*f D*) is the change in transmission caused by the detection of infections and it is defined as *ϵ* times *f*_*n*_*D*_*n*_ *− f*_*o*_*D*_*o*_. The other parameters are as described above. Note that if we relaxed the assumption that *ϵ* was unchanged by evolution, the numerator on the left hand side would have instead been Δ(*ϵf D*). Regardless, this inequality states that a more harmful variant would be favored by evolution if the percent decrease in ineffective control is less than the percent increase in the transmission rate in the absence of detection. Notice the parallels with the analogous result using the classic virulence-transmission trade-off, where the key inequality was that the percent decrease in survival must be less than the percent increase in transmission. Hosts that survive continue to transmit analogously to how hosts with undetected infections or ineffective interventions continue to transmit.

Nevertheless, there is a key difference between Inequalities 6 and 13. I previously showed using an example that host mortality is unlikely to constrain the evolution of pathogen virulence for pathogens with initially low infection fatality rates *F*_*o*_. Again consider the same hypothetical pathogen with an infection duration of 5 days and an *R*_0_ of 5, but this time, focus on the cost of detection. Assume individuals can be infectious 1 day before developing symptoms and 4 days after. Also, assume that an individual becomes less likely to attend school, work, or other social events with increasingly severe symptoms, and that the vast majority of transmission occurs during these activities. Using this information, we can ask under what circumstances a new variant that causes the average person to stay home one day would be able to displace a less severe variant that causes the average person to not stay home at all (i.e. virtually no initial cost of virulence as in the mortality cost example above). Using the above details, we can calculate the key parameters: *ϵ* = 4*/*5, *f*_*n*_*D*_*n*_ = 1*/*4, *f*_*o*_*D*_*o*_ = 0. Plugging these values into Inequality 13 leads to the conclusion that this variant would only be able to invade if it were accompanied by a 20% or larger increase in transmission. This can be visualized in Fig. 3 if the x-axis label was changed from “infection fatality rate (*F*)” to “the reduction in transmission due to detection (*ϵf D*)”. If we were to put this example with detection costs on the same scale as the previous example with mortality costs, transitioning from a 0% infection fatality rate to a 1% infection fatality rate is an equivalent cost to transitioning from a 0% chance of staying home for one day to a 5% chance. While the former would almost certainly be documented if it were to evolve in human populations, the latter almost certainly would not. Again, these changes would only be evolutionarily favored if they led to an increase in transmission of approximately 1% of more (Inequalities 6 and 13).

Presumably many infectious diseases, including non-human diseases, could be constrained by costs of detection. However, detection would not have much impact on limiting disease severity if large fractions of the infectious period occurred prior to the time when detection would be possible (i.e. *ϵ* is small), if reductions in transmission were small following the detection of infection (i.e. *f*_*o*_ is small) or if a very small fraction of infections were detected (i.e. *D*_*o*_ is small). Likewise, host-induced mortality can be a strong constraint on virulence evolution if the infection fatality rate *F*_*o*_ is large. To determine whether virulence is more strongly shaped by a mortality-transmission trade-off or a detection-transmission trade-off, one can combine Inequalities 6 and 13 to ask:

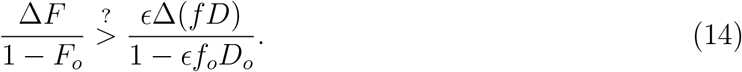

When the left-hand side of the above expression is larger than the right-hand side, mortality will impose a stronger constraint on virulence evolution than detection, and vice versa. For a pathogen with low virulence, the denominators on both sides are close to one meaning that we can visualize this inequality using only the numerators (Fig. 4). This demonstrates that for pathogens with relatively low virulence, detection will generally be a stronger constraint on virulence evolution than mortality.

**Figure 4:**
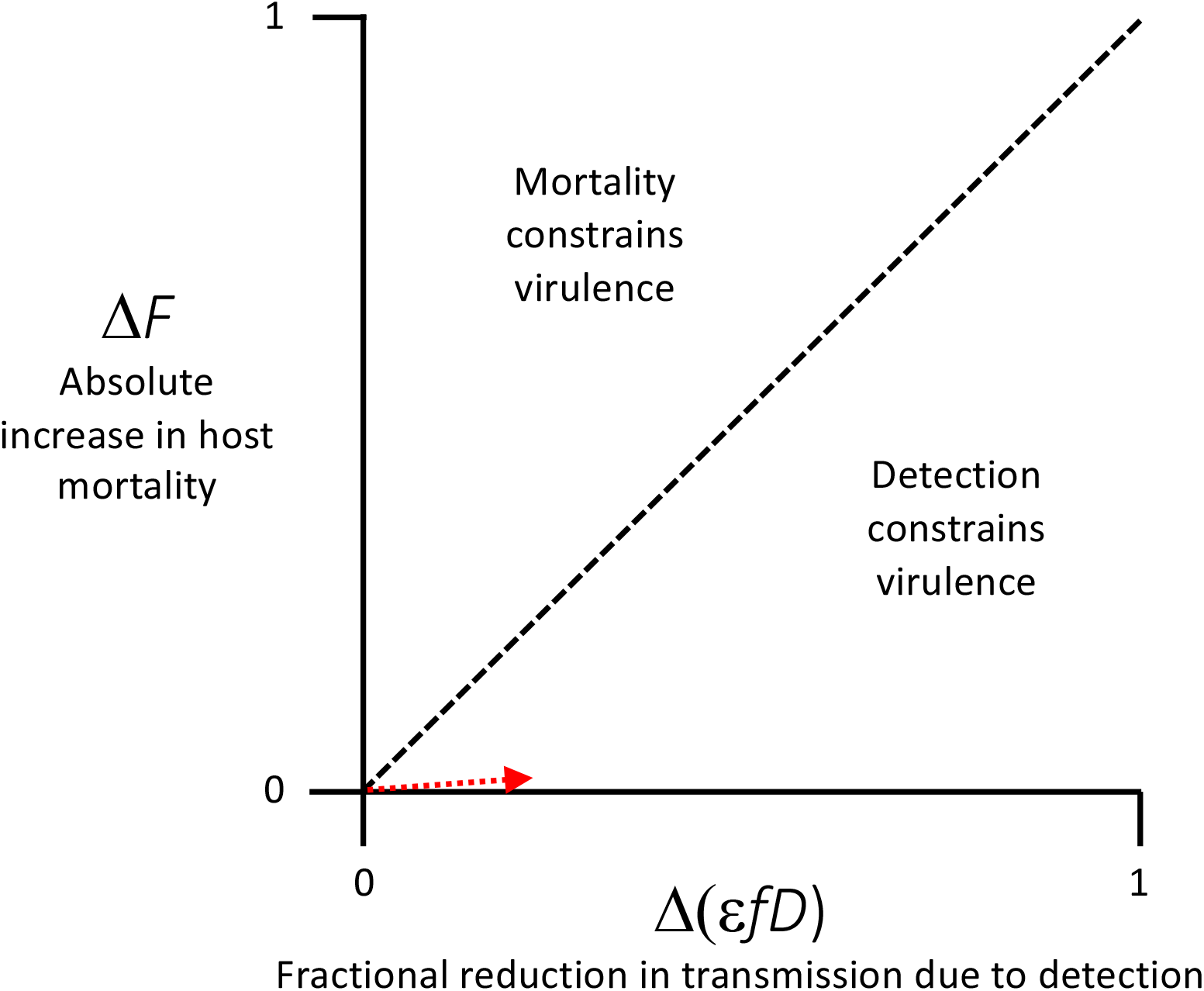
A graphical representation of Inequality 14 for a pathogen with initially low virulence. The cost from an x% reduction in average transmission is equivalent to the cost from an x% increase in the infection fatality rate. The dashed line is the 1:1 line. Above the dashed line, virulence is constrained by mortality costs, and below it, detection costs. The dotted red arrow depicts the example provided in the main text of a pathogen with initially low virulence that evolves higher virulence in the form of either killing 1% of infected hosts (i.e. roughly equivalent to SARS-CoV-2 infection) or causing infected hosts to stay home for one day (roughly equivalent to infection with a virus that causes the flu or the common cold). Notably, the cost of the former is much smaller than the cost of the latter despite the fact that most would consider the former more virulent than the latter.

## Discussion

The new conventional wisdom says that pathogens evolve to balance the cost and benefits of virulence and its associated traits. Typically, the cost of virulence is assumed to be a truncated infectious period due to disease-induced host mortality. Here I have argued that this cost is too weak in most systems to constrain virulence. To do this, I have rewritten the virulence-transmission trade-off equation in terms of infection fatality rate (i.e. the fraction of infections that result in host death *F*) rather than in terms of the per day infection-induced death rate (i.e. *α*). This formulation makes explicit that mortality-based evolution of virulence theory predicts that a novel variant would be able to displace an existing pathogen variant if the percent decrease in host survival is less than the percent increase in the rate of transmission (Inequality 6). However, this is an extremely weak evolutionary force for all but the most virulent pathogens (Figs. 2 and 3). I instead propose that the main cost of virulence is due to behavioral changes that result from the detection of infection. Using a modified SIR model that explicitly allows for costs of detection, I show that a detection cost can be a much stronger constraint on virulence evolution than host-induced mortality (Fig. 4).

It has previously been noted that costs of mortality are small (Ebert and Bull 2003; Bull and Lauring 2014). I provide an analytical expression for precisely how small (Inequality 6). This expression states that when infection fatality rates are low, a percent change in transmission rate can balance an equivalent absolute change in the infection fatality rate (Figs. 2 and 3). Under the assumption that mortality limits disease severity, a 1% increase in transmission can thus justify something as harmless as a virus that causes the common cold evolving to become something as deadly as SARS-CoV-2, yet a common cold virus has never evolved to be so deadly. Certainly something else must constrain virulence.

Host behavior, unlike host mortality, is widely recognized to influence infectious disease dynamics (Funk et al. 2010; Stockmaier et al. 2021). It only stands to reason then, that when infection-induced behavioral changes affect opportunities for onward transmission, and disease severity alters the degree of behavioral change, there will be opportunities for natural selection to shape disease severity. This idea was originally made by Ewald (1983, 1994) to argue that vector transmitted diseases should evolve to be more virulent than directly transmitted diseases since they do not rely on their hosts for dispersal. However, definitive data supporting Ewald’s argument regarding transmission mode are still lacking (Leggett et al. 2017). While a disconnect between his assumptions and his conclusions may be due to variation in system specific details (Day 2002a), support for mortality costs in the case of myxomatosis (Fenner 1983; Dwyer et al. 1990) led to mortality being viewed as a reasonable constraint on virulence evolution. I have shown that this is not the case when infection fatality rates are low, and it may explain why surprisingly few data support the idea that mortality acts as a constraint on pathogen evolution (Bull 1994; Lipsitch and Moxon 1997; Ebert and Bull 2003; Alizon et al. 2009; Bull and Lauring 2014; Cressler et al. 2016; Acevedo et al. 2019).

At some level, this conclusion may be obvious. Both mortality and detection could in principle constrain virulence evolution, but death is usually a rare outcome of infection whereas detection is usually a common outcome. It thus follows that detection costs may often be larger than mortality costs.

Few data are yet available to quantify precisely how large detection costs are, but the data that do exist suggest these costs are quite large in comparison to fully asymptomatic infections. A study of influenza-like-illness (ILI) during the 2009 influenza pandemic found that people with ILI reduced their per day contacts by 75%, and the average duration of contact also declined (Van Kerckhove et al. 2013). Despite this reduction, two thirds of transmission was attributable to symptomatic infection, suggesting a steep trade-off between contact rate and infectiousness given contact (Van Kerckhove et al. 2013). Another study using seasonal influenza documented a negative correlation between morbidity scores and activity levels among people with detected infections, and even proposed that this reduction in activity may pose a constraint on virulence evolution in that system (McKay et al. 2020). Similarly, a survey study on behavioral change following diagnosis with various sexually transmitted diseases reported that 71% of men modified their behavior (e.g. increased condom use, reduced frequency of sex, etc.) following diagnosis. Isolation and quarantine following the detection of SARS-CoV-2 infection in an individual or in a close contact of an individual likewise is thought to have large impacts on disease transmission (Keeling et al. 2020). Nevertheless, more data are still needed to establish whether the magnitude of these detection costs are typical for human diseases.

Similar magnitude effects are seen in the animal world. When wild house mice were experimentally injected with lipopolysaccharide (LPS) to induce disease symptoms, 40% of the mice disconnected entirely from their social groups (Lopes et al. 2016). Although these mice did not have an infectious disease, the change in behavior brought on by a general immune response would have substantially reduced opportunities for pathogen transmission if it were brought on by a pathogen (Lopes et al. 2016). Vampire bats injected with LPS also showed large changes in behavior, with 85% less time spent grooming conspecifics and 19% less time spend being groomed by conspecifics (Stockmaier et al. 2020). Analogous patterns were found in guppies infected with an ectoparasite. Guppies typically form groups called shoals, but when infected guppies were added to otherwise healthy populations, the healthy fish actively avoided the infected guppies causing fission events at twice the rate of controls, and associations when they did occur were half as long in duration (Croft et al. 2011).

In a eusocial ant species, when colony workers were experimentally infected with a fungal pathogen, the social network of the colony changed in ways that reduced opportunities for disease transmission, including a shift such that experimentally infected worker ants spent 20% more time outside of the nest than controls (Stroeymeyt et al. 2018).

Notably, detection costs may even be playing a role in limiting virulence for some of the systems where virulence-transmission trade-offs have been best documented. For example, in *Mycoplasma gallisepticum* where prior immune history enhances the spread of highly virulent strains (Fleming-Davies et al. 2018), interaction rates between birds are approximately 15% lower for infected birds than non-infected birds (Faustino et al. 2004). Likewise, for monarch butterflies infected with the parasite *Ophryocystis elektroscirrha*, reductions in mating success that prevent transmission to offspring actually provide a stronger constraint on parasite load than mortality, captured as pupal emergence (De Roode et al. 2008), although perhaps not significantly so.

While the above data may be subject to some of the same publication biases that have previously plagued trade-off theory (Acevedo et al. 2019), the effects in the above studies tend to be highly significant and a mechanistic basis for the effects seem logical (Ewald 1983, 1994). Moreover, there is a long history of humans altering their behavior in response to the detection of infectious disease (Curtis 2014).

Admittedly, the model described in Eqs. 7-11 is more complicated than absolutely necessary. Very similar conclusions could have been derived from a standard SIR model where the transmission rate *β* is separated into two components, the rate of contact and the probability of infection given contact. In that model, if the rate of contact declines with increasing virulence, that cost could constrain virulence evolution (Day 2001, 2002a). However, it is difficult to intuit the reasonableness of changing these parameters since they are composite parameters like transmission rate itself. By introducing the biologically meaningful parameters *ϵ, f*, and *D*, I hope to have provided a clearer framework for thinking about the basis of detection costs.

Along similar lines, I have followed the standard SIR model assumption that mortality risk is constant for the duration of an infection. This is typically not true, with mortality typically occurring towards the later phase of infection. If this were incorporated into my analysis, the effect is that the cost of mortality would be even weaker than I have calculated, further strengthening my claim that mortality costs are typically too weak to constrain virulence evolution.

Note that despite the use of the term “detection cost”, my argument is agnostic as to the exact mechanism causing the change in interactions. Multiple mechanisms can result in reduced transmission, and have been documented in human and non-human hosts. Detected infections can result in reduced transmission if infected hosts are too ill to go about their normal routine and thus contact fewer susceptible hosts (e.g. Stockmaier et al. 2020), if they take action to avoid spreading an infection through intentional behavioral modification (e.g. Süss et al. 2011; Stockmaier et al. 2021), if they seek treatment to end infection earlier (e.g. Alizon 2020), or even if they are avoided by others who notice that they are ill (e.g. Curtis 2014)).

Perhaps the greatest challenge moving forward is to test this theory experimentally. The difficulty of doing so stems from being able to create conditions that are close enough to field conditions such that they allow for changes in behavior that limit transmission following the detection of infection. Such laboratory experiments may prove too difficult to design, and may ultimately mean that tests of this theory must be performed in the field.

It is worth noting that virulence has been defined differently by different researcher (Read 1994; Thomas and Elkinton 2004; Alizon and Michalakis 2015; Cressler et al. 2016). For example, virulence can be defined as the pathogen induced reduction in host fitness (Read et al. 2015), as the per day pathogen-induced host mortality rate (Anderson and May 1982), as the fraction of hosts that die from infection (Day 2002b), or in numerous other ways (Thomas and Elkinton 2004; Cressler et al. 2016), and these differences can lead to fundamentally different conclusions (Day 2002b). Here I have defined virulence as the severity of disease signs or symptoms caused by infection with a pathogen. The argument that I have put forward applies to this definition of virulence specifically. While it may apply to other definitions of virulence as well, this application relies on correlations in “virulence scores” between the definitions.

Despite my above argument, there are some situations in which a mortality cost can provide a stronger constraint on the evolution of virulence than a detection cost (Inequality 14, Fig. 4). For example, mortality costs appear to have been major drivers of pathogen evolution for myxomatosis (Fenner 1983), Marek’s disease virus (Read et al. 2015), and some bacteriophages (Messenger et al. 1999). Notably, accounting for pathogen-induced host mortality is important for accurately modeling disease dynamics in these systems (Dwyer et al. 1990; Berngruber et al. 2013; Atkins et al. 2013).

Here I have assumed that the benefits of virulence come from a correlation with transmission rate (Fig. 1). As shown by Inequality 14, the precise benefit of virulence does not impact whether virulence is more strongly constrained by mortality or detection. Numerous alternative theories have been proposed to explain why pathogens maintain virulence even in cases where virulence itself is not obviously beneficial (Frank 1996; Alizon and Michalakis 2015; Cressler et al. 2016). Some of these theories include that multilevel selection leads to the evolution of virulence levels that are non-optimal at the between-host scale (e.g. Levin and Bull 1994; Mideo et al. 2008), that spatial structure imposes dispersal or persistence costs of high virulence (e.g. Boots et al. 2004), that environmental feedbacks limit the relationship between the basic reproductive number *R*_0_ and optimal virulence (Lion and Metz 2018), that virulence is not adaptive in the context where it is being studied but adaptive in another context (Ebert 1999), that bottlenecks prevent the evolution of optimal virulence (Bergstrom et al. 1999), or simply that there is no heritable variation for virulence on which selection can act. Notably, these theories have largely been developed to explain why trade-off theory seems not to apply generally, and while the theoretical impacts of these factors have been demonstrated, the extent to which they play out in the real world is still unknown. Assuming detection is a main factor in limiting the evolution of virulence, it begs the question of how this knowledge might be used. Ebert and Bull (2003) previously argued that virulence management is not practical when it relies on indirect selection using trade-off theory. They instead proposed that efforts would be better aimed towards selecting against virulence directly. I propose that if virulence is truly constrained by a cost of detection, then efforts to increase detection would leverage trade-off theory while also directly selecting against virulence, provided the increased detection effort maintains the correlation between detection and virulence. Likewise, as we saw during the early days of the COVID-19 pandemic, surveillance programs are often designed to catch clusters of symptomatic infection (Kerr et al. 2021). This may unintentionally provide additional evolutionary benefits in that more virulent pathogens will be more likely to be caught and stopped.

## Acknowledgments

I thank T. Day, G. Dwyer, and A. Read for comments on previous versions of the text. This work was supported by National Science Foundation grant DEB-1754692. The funders had no role in study design, data collection and analysis, decision to publish, or preparation of the manuscript.

## Supplemental Information

### Classic virulence-transmission trade-off

Under what conditions will a new mutation *n* be able to invade and displace an original pathogen variant *o*. Assuming Eqs. 1-3, a new mutation can invade when:

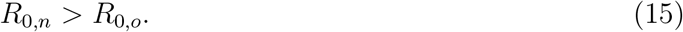

Substituting equation 4 into inequality 5 yields

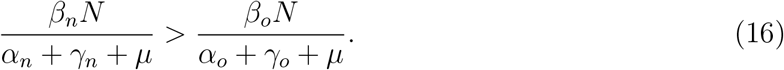

Multiplying both sides by *α*_*o*_ + *γ*_*o*_ + *µ* and dividing both sides by *β*_*n*_*N* yields

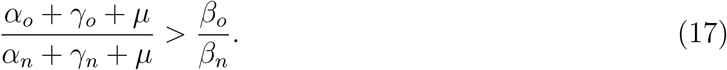

Multiplying both sides by negative 1 and then adding 1 to each side yields

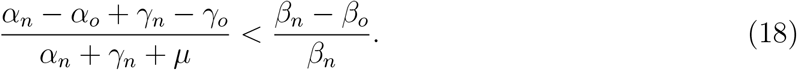

Under the assumptions of a virulence-transmission trade-off constrained by disease-induced mortality, *γ*_*o*_ = *γ*_*n*_ and so the above can be rewritten

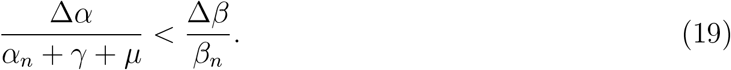

where Δ*α ≡ α*_*n*_ *− α*_*o*_ and Δ*β ≡ β*_*n*_ *− β*_*o*_.

### Rewriting the virulence-transmission trade-off in terms of *F*

In the model described by Eqs. 1-3, there are two ways an infected host can leave the infected class *I*: 1) through death caused by infection and 2) through recovery. Since both the infection-induced mortality rate *α* and the recovery rate *γ* are assumed to be constants over time in the standard SIR model, the fraction of hosts that leave the infected class through infection-induced death is simply described by the equation (Day (2002b) previously presented the generalization where rates are variable over time):

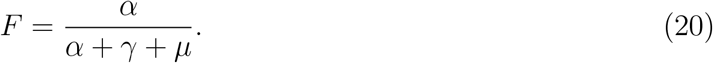

I use the above equation to solve for *α*_*o*_ in terms of *F*_*o*_ and *γ*_*o*_, which yields

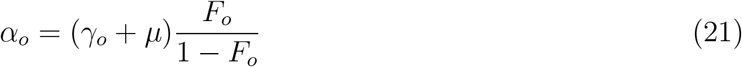

Returning to eq. 19, substituting in eq. 21 for *α*_*o*_ and the equivalent for *α*_*n*_, and then simplifying gives

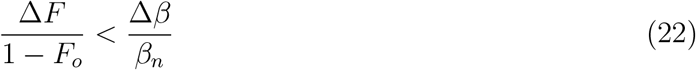

where Δ*F ≡ F*_*n*_ *− F*_*o*_ and Δ*β ≡ β*_*n*_ *− β*_*o*_. Inequaliity 22 thus shows that a new mutation will be favored if the percent change in the infection survival rate is less than the percent change in the transmission rate.

### Detection costs

Starting from Eq. 12, a new mutation can invade when:

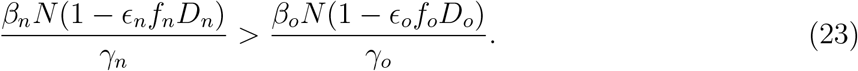

Multiplying both sides by *γ*_*o*_ and dividing both sides by *β*_*o*_*N* (1 *− ϵ*_*o*_*f*_*o*_*D*_*o*_) yields:

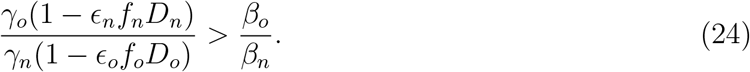

Again multiplying both sides by negative 1 and adding 1 to each side yields:

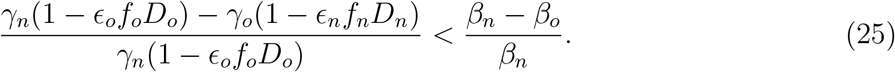

Assuming *γ*_*n*_ = *γ*_*o*_, we arrive at our final solution:

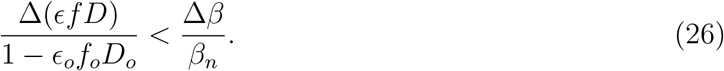

